# Low-dimensional olfactory signatures of fruit ripening and fermentation

**DOI:** 10.1101/2024.06.16.599229

**Authors:** Yuansheng Zhou, Thomas F. O’Connell, Majid Ghaninia, Brian H. Smith, Elizabeth J. Hong, Tatyana O. Sharpee

## Abstract

Odors provide an important communication channel between plants and animals. Fruits, vital nutrient sources for animals, emit a complex array of monomolecular volatiles. Animals can use the structure of these mixtures to assess properties of fruit predictive of their nutritive and reproductive value. We analyzed the statistics of fruit odor mixtures sampled across stages of ripening and fermentation to find that they fall on a low-dimensional hyperbolic map. Hyperbolic maps, with their negative curvature and an exponentially expanding state options, are adept at describing hierarchical relationships in the data such as those arising from metabolic processes within fruits. In the hyperbolic map, samples followed a striking spiral trajectory. The spiral initiated near the map’s core, representing the under-ripe phase with specific profiles of monomolecular volatiles. Progressively mapping along the unfolding spiral trajectory were scent mixtures corresponding to ripening, and then rotting or fermentation. The unfolding process depended on the specific fermentation processes that dominated in the samples, determined largely by the microbes (e.g. bacteria or yeast) present in the sample. These results generalized across fruit types and describe trajectories in the natural odorant space with significant behavioral relevance for insects.

## Introduction

Understanding chemical communication is critical to understanding the complex networks of interactions between organisms forming any ecosystem. The vast majority of organisms on the planet use chemical communication to coordinate the core behaviors necessary for survival and reproduction, which can include identifying suitable habitats, locating sources of energy and nutrients, finding mates, and avoiding danger^1-3^. However, the odor stimulus space is large and high-dimensional, posing unique challenges for understanding olfaction. In other sensory modalities, sensory stimuli are easily organized along physical axes like wavelength of light or frequency of sound, and these axes are reflected in the topographical organization of the corresponding sensory code in the brain^4,5^. In contrast, the principal organizational axes of odor space are not obvious. The protein receptor repertoires that detect odorant molecules are correspondingly large and diverse^6,7^, and attempts to organize odors along axes defined by their physicochemical or structural properties, such as size, hydrophobicity, or functional groups, only partially predict the perceptual relationship between odors^8-10^. These challenges motivate additional approaches to developing maps of olfactory stimulus space.

Here, we investigated the idea that odor space can be efficiently organized around the relationships among odors with respect to their occurrence in natural odor sources of behavioral significance to animals. In the natural world in which animal brains evolved, specific volatiles emanating from an odor source are generated by biochemical and metabolic processes in the natural source and its associated microbes^11-15^. The abundance and co-occurrence of specific compounds is distinct for different odor sources and signals important information about the source, such as its suitability as an energy source or the potential presence of toxins^16-19^. Thus, the odor space of the natural world is highly structured, and this structure likely carries behavioral meaning to animals. We refer to the complex mixtures of monomolecular volatiles emanating from naturally occurring odor sources as natural mixtures. We hypothesized that investigating the properties of natural mixtures, in particular, identifying low-dimensional descriptions of the ethologically relevant variation between them, would yield a useful organizational framework for olfactory stimulus space.

Specifically, we considered the odor space of fruits in the natural world. Fruits of flowering plants are nutrient rich structures that form from the ovaries of flowers and are designed to protect seeds and enhance seed dispersal^20^. Fruits, and their associated commensal microbes, attract a diverse set of animals, including birds, mammals, and insects, that consume the fruits as rich sources of carbohydrates, proteins, vitamins, and other essential nutrients^21-23^. Additionally, insects such as fruit flies may be attracted to fruits via odor cues as productive sites for egg laying^24,25^. Insect larvae derive nutrients from the fruit pulp and in some cases from the yeast that grow on fruit, a major source of protein for developing larvae. Ripeness state, and microbial colonization by bacteria and/or yeast, provide important cues to animals about the suitability of the source for ingestion or for supporting reproduction^26^. Humans prefer ripe fruits with high sugar content^27,28^ and avoid spoiled fruits, while the scent of alcohol may signal to many mammals the presence of ripe, fermenting, nutritious fruit^29,30^. Different fruit fly species may prefer cues preferentially associated with unripe, ripe or fermenting states to avoid inter-species competition^18,31^ or to match the demands of their current physiological state^32^. Fruits undergoing yeast fermentation are of particular significance to female and larval fruit flies, which feed on yeast as an important source of protein.

Like other plant odors, such as floral odors^33^, fruit odors are typically comprised of many different monomolecular compounds^34^. Previous studies of ripe fruits from different strawberry varieties, for example, identified dozens of different chemical compounds across varieties^27,28^.

The differences in chemical composition were correlated with variation in the palatability and attractiveness of the fruit to human subjects^27,28^. Other studies designed to evaluate ripeness states for fruit fly preferences have shown that different species prefer different states^18^, and that the odor preferences of those species can be explained by key odorants that are given off at those ripeness states by the fruits or from green leafy volatiles. These studies have either implied indirectly or via direct bioassay that fruit and vegetative odors are important for detection and identification of fruits in different states.

Thus, an understanding of the physical space of odors that fruits emit, and in particular the statistical relationships among those ligands as the fruits change state, is a fundamental step towards understanding the behavioral value signaled by these odors. We argue this is an important step toward understanding the neural mechanisms that mediate odor-driven behavior. Analogously, investigations of the statistics of naturalistic visual and auditory scenes have led to significant insights into the structure of sensory codes for those modalities^35,36^. The goal of the current study was to examine the statistical relationships of volatile compounds in natural odor mixtures as they relate to the ecological significance of these odors. We analyzed odorants emitted by five different types of fruits (strawberries, banana, mango, kiwi, and peach), across different ripeness states and different types of fermentation. We show that a curved (hyperbolic) model provides a better fit for this space than linear methods, similar to floral odors^37^, and identify paths or trajectories in odor space associated with distinct metabolic states with differing ecological value. We discuss the implications of a curved odor space for understanding the origins of its structure in the metabolic pathways that produce natural odors.

## Results

### Chemistry of fruit odors

We analyzed two independently collected data sets designed to sample different fruits in different stages of ripening and fermentation (Fig 1 with supplements), which we expect to have varying behavioral value to animals. The first data set measured ripening, whereby ‘green’ unripe or ‘ripe’ whole strawberry fruits were allowed to sit at room temperature as they went through visually confirmed ripening, classified as ‘green/blush/unripe’, ‘red/ripe’ or ‘brown/overripe’. The second data set used five different fruits – banana, kiwi, mango, peach and strawberry – which were homogenized, introduced into a vessel sealed with a fermentation airlock, and allowed to proceed through two different types of fermentation. Approximately half of the samples of each type of fruit proceeded through ‘uncontrolled’ fermentation at room temperature, driven by the microbes endogenous to the fruits as purchased from the grocer, similar to the strawberry fruit in the first data set. The other half of the homogenized fruit samples were chemically sterilized and inoculated with wine yeast, which then dominated the fermentation process.

**Figure 1:**
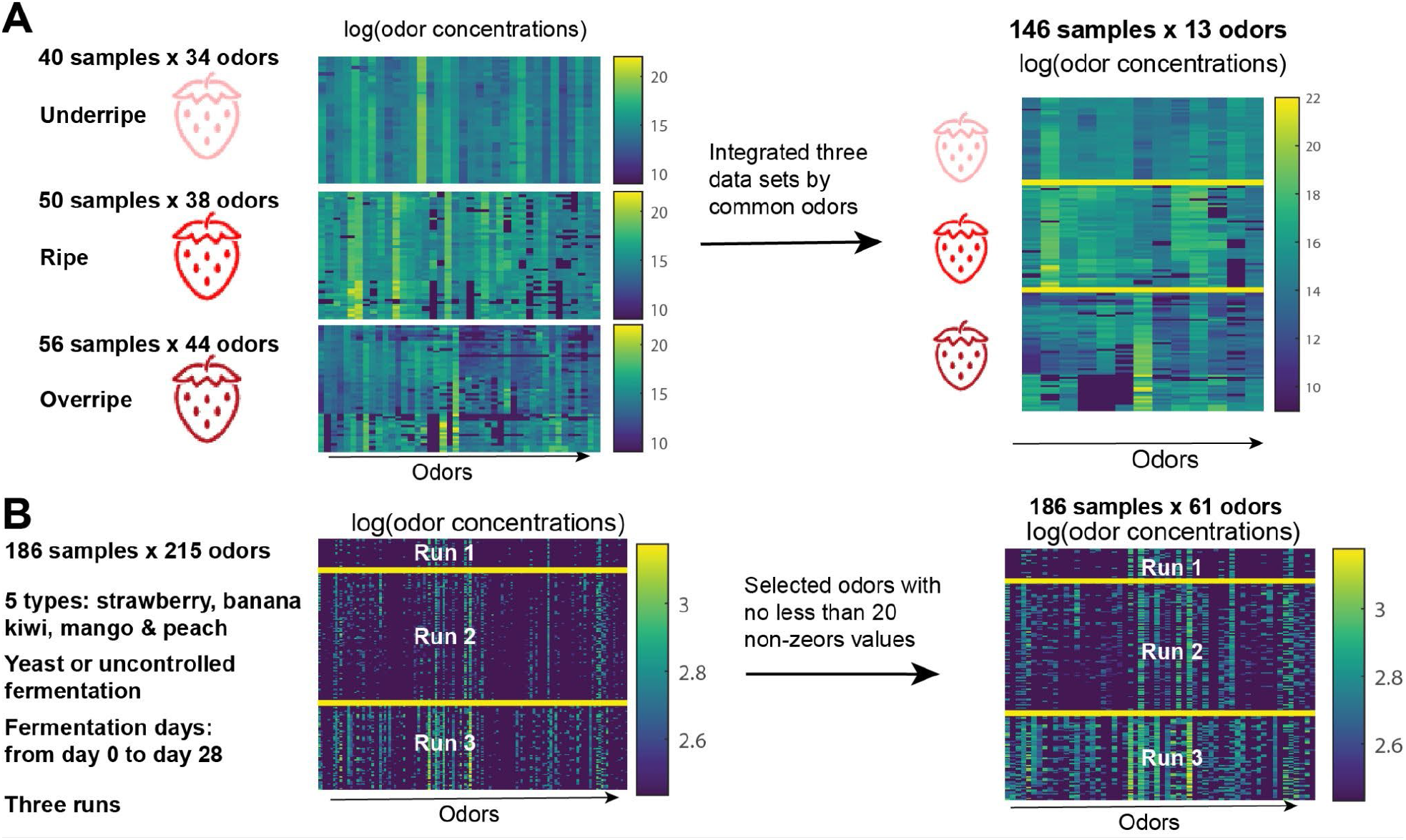
Processing of strawberry and fermentation data before low dimensional embedding. (A) The strawberry data contains under-ripe, ripe and overripe data sets which share 13 common odors. The three data sets are integrated by selecting the common odors. (B) The fermentation data is sparse, and therefore we omitted odors with less than 20 non-zero values. The concentration values in both datasets were log-transformed before doing low dimensional embedding.

We used gas chromatography-mass spectrometry (GC-MS) to analyze the chemical volatile profile of the headspace of these fruit samples in different ripeness or fermentation states. For the whole strawberry dataset, dozens of samples (individual strawberry fruit) from each ripening state (unripe, ripe, or overripe) were analyzed using dynamic headspace analysis (Fig 1 supplement 1). For the fruit homogenate dataset, the headspace above the odor source in the fermentation vessel was sampled at 0, 1, 2, 3, 5, 11, or 14 days of fermentation using static headspace analysis by solid phase micro-extraction (SPME) coupled to GC-MS; three independent fermentation runs were analyzed for banana, mango, and kiwi (Fig 1 supplement 2). Some fermentation runs were additionally analyzed at 21 and 28 days of fermentation. We identified dozens of monomolecular volatiles emanating from each source, some of which were common across all states and others characteristic of just one or two states (Fig 1 supplement 3). Several chemicals identified from ripe strawberries were also identified in previous analyses of ripe strawberries^27,28^. Thirteen compounds (Fig 1 supplements 1 and 2, rows 1 through 13) were common to all ripeness states. We focus our dimensionality reduction on this common set of monomolecular odorants. Other compounds were characteristic of one or two states but not all three, and are summarized in Supplemental Table 1.

In the individual strawberry fruit dataset, samples from green fruits, or from early ripe fruits (e.g. day 0,1) were fairly uniform with respect to composition and, to some extent, the abundance of specific compounds. With progressive ripening, an overall increase in sample-to-sample (columns) variation was observed (Fig 1-Supplement 1). Changes across ripeness states occur in part because of changes in metabolic pathways in fruit cells that produce these volatiles. However, these changes could also arise because of biochemical pathways, like sugar fermentation pathways, that become dominant as the number of microbes in the sample increase. For example, several of the compounds we identified are also common to yeast (Fig 1 supplement 3), and thus these chemicals could be from the fruit and/or yeast presence during fermentation. Finally, sample-to-sample variation within ripeness state could arise because of variation in individual fruit states as they move through ripening and fermentation. For example, some fruits may be entering fermentation but still be visually classified as ripe.

Several chemicals we identified in one or both data sets have not been identified in other studies of strawberry fruits (blue chemical names in Fig 1 supplement 3). This result partly stems from the paucity of studies of unripe and fermented fruits, which we included here because of the relevance of those states to humans and flies^28,38^ For our analyses of the odor space of fruits, we focused on a list of 13 chemicals which were observed across all states in our samples with high confidence, and also identified in two other studies of strawberry fruits^27,28^.

Notably, styrene was detected in all sample sets, yet it was not reported in other analyses of, for example, strawberry fruits^27,28^. Note that previous analyses of styrene emphasized only samples of ripe fruits, whereas we includes advanced fermentations states relevant to insects. Styrene can have a sweet, balsam, floral or plastic odor to humans. In our samples, it appeared more frequently and in larger concentrations in later stages of fermentation of all types of fruits. In fact, it was particularly important in determining the spiral shape of trajectories through fermentation states. Styrene is a natural product that has been reported to occur at low levels in some foods, including strawberry and pear fruit^39,40^ and in samples of yeast and fungal spoilage^41,42^. We suspect that our samples contained styrene because of the emphasis on sampling more advanced fermentation states that may be of relevance to fruit flies. Therefore, it might be a natural product in our samples, and in particular it might arise from fungi and bacteria in later stages of fermentation.

Nevertheless, styrene could also occur as a contaminant from exposure to plastics (polystyrene) during harvesting, storage and shipment of fruits. In the absence of proof that it is or is not a natural product in our samples, we analyzed data both with and without styrene, and the hyperbolic nature of the remaining fruit space was still clear and robust.

### Hyperbolic geometry of fruit ripeness and fermentation process

We proceeded to analyze global geometry properties of the two datasets of odorants: the combined strawberry dataset, cf. Fig.1A, consisting of 146 samples across the three fermentation stages (Fig. 1-Supplement 1; Supplementary Table 1) and the fermentation dataset consisting of five types of fruits, two fermentation conditions, 1-3 runs, and progressive sampling at successive fermentation days (Fig. 1B; Fig. 1-Supplement 2; Supplementary Table 2). In the latter, we selected 61 odors with more than 19 non-zero concentration values across all samples (Fig.1B). The odor concentrations were log-transformed before downstream analysis.

We first analyzed the geometry of strawberry and fermentation datasets using methods based on non-metric multidimensional scaling that we previously developed^43^. Briefly, the method adjusts the curvature (or size in units of inverse curvature) of the embedding space to achieve a linear Shepard diagram between original and embedded distances^43^. We observed that hyperbolic space provided a better fit than the Euclidean space (Fig. 2). For example, the pairwise distance plots show a nonlinear relationship in 3D Euclidean multi-dimensional scaling (MDS) embedding (Fig. 2C, E), but a linear relationship in 3D hyperbolic embedding (Fig. 2D, F), for both the whole fruit and fruit homogenate dataset.

**Figure 2:**
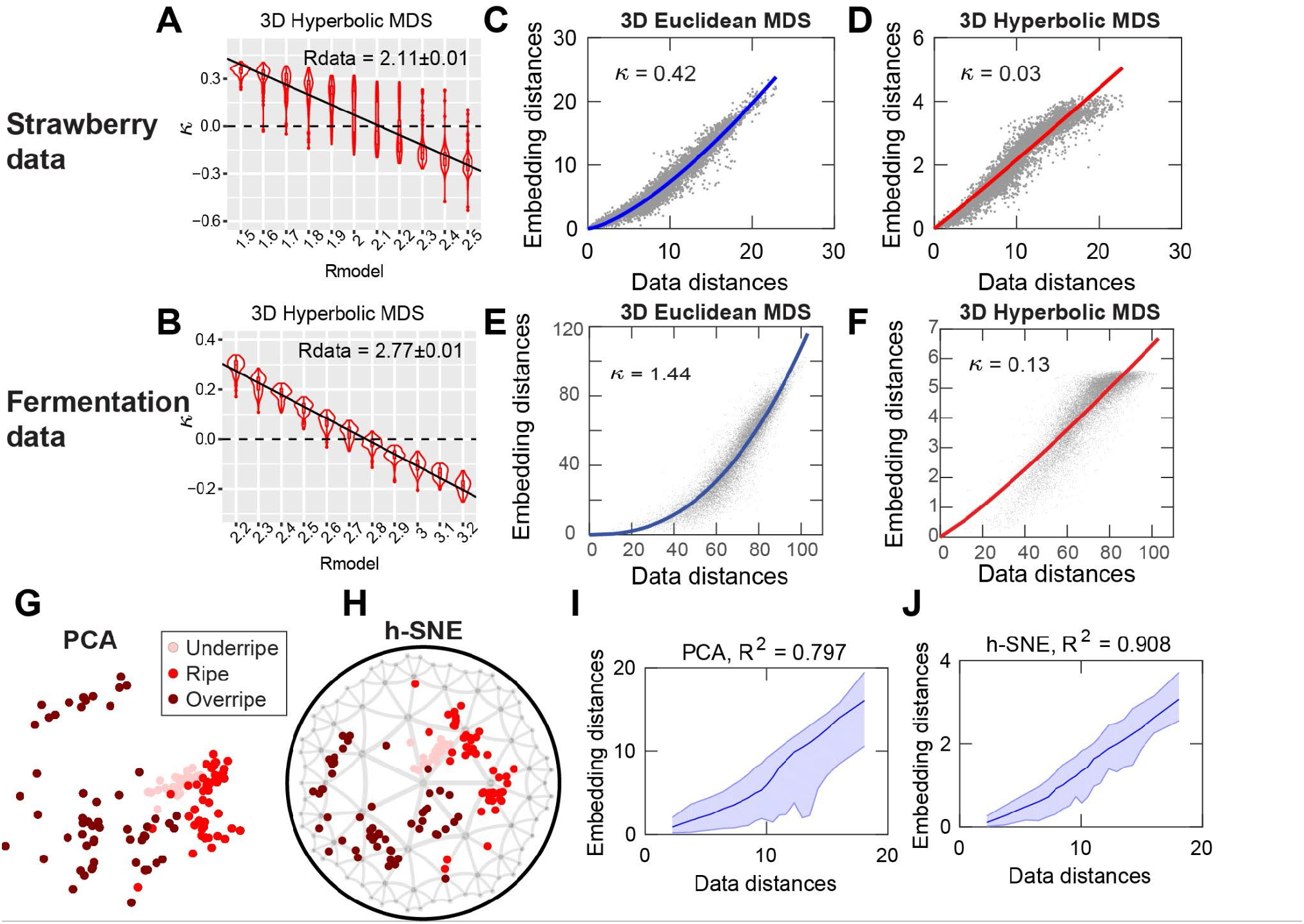
Hyperbolic geometry describes strawberry and fermentation data. We performed hyperbolic multi-dimensional scaling (HMDS) with a different *R*_*model*_ for the strawberry (**A**) and fermentation (**B**) datasets, plotting the curvature of Shepard diagram fitting versus *R*_*model*_. The violin plots show the curvature statistics from 100 repetitions of the embeddings. The details of the method were previously described^43^. (**C**) 3D pairwise distances plots of 3D Euclidean embedding of strawberry data; the curvature index of the fitting function *y* = *αx*^*k*+1^ is indicated in the panel. (**D**) 3D hyperbolic embedding of strawberry data with *R*_*model*_= *R*_*data*_= 2.11. (**E**) 3D Euclidean embedding of fermentation data. **(F)** 3D Hyperbolic embedding of fermentation data with *R*_*model*_ = *R*_*data*_ = 2.77. (**G**). Visualization of strawberry data using the first two components of PCA. The pink, red and dark red colors represent under-ripe, ripe and overripe strawberry samples. (**H**) 2D h-SNE embedding of strawberry data. (**I**) The distance plot of PCA embedding. (**J**) The distance plot of h-SNE embedding.

To further evaluate Euclidean embeddings, we performed PCA for strawberry data and embeded the samples by the first two principal components (Fig. 2G) and observed a nonlinear relationship between data distances and embedding distances (Fig. 2I). These results show that PCA embedding in Euclidean geometry is not appropriate for the hyperbolic strawberry dataset. Hence, we applied hyperbolic t-SNE (h-SNE), previously developed^43^ to visualize the data. The h-SNE gives a clear spiral shape of a point cloud which starts from the original point (Fig. 2H). The distance plot also shows a better data distance preservation with *R*^2^ = 0.908, compared to *R*^2^ = 0.797 in PCA embedding (Fig. 2J).

When embedded into 3D hyperbolic space, both datasets were best described by maps with approximately the same hyperbolic radius: *R*_*data*_ = 2.11 for strawberry data (Fig. 2A), and *R*_*data*_ = 2.77 for fermentation data (Fig. 2B). The size of the hyperbolic map can be used as a measure of the hierarchical depth of the dataset^43^. The radius is larger for fermentation data, because it is more complex than strawberry data. The fermentation data contains multiple fruit types, and the higher complexity can be reflected in the larger hyperbolic radius of the embedding^43^.

### Spiral progression of strawberry ripeness

To visualize trajectories in the reduced space that may be related to behavioral meaning or value, we performed hyperbolic t-SNE embedding of the data^43^. We first analyzed the whole strawberry dataset where the method showed that the under-ripe, ripe and over-ripe strawberries follow a spiral trajectory (Fig. 2H). The simplest spiral is one-parameter Archimedean spiral in which the radius is proportional to the angle:

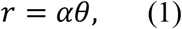

where *α* describes how fast the spiral expands with the angle. To fit the embedding points with the Archimedean spiral, we performed distance-invariant transformation by performing reflection and rotation which were optimized for the fitting. The optimal fitting parameter was *a* = 0.32 (Fig. 3B), and the relationship between radii and angles are shown in Fig. 3C. The spiral length of a point (*r*_*i*_, *α*_*i*_) is defined as:

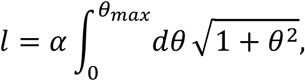

where *α*_*max*_ = *α*_*i*_ and *a* is the spiral coefficient in Eq. (1). To examine how variation in fruit odor mixtures changed with ripeness stage, we performed hyperbolic t-SNE in 3D and found the optimal 2D projection disk that gives the best spiral fitting of projected points in the disk (Fig. 3D). The parameter for the optimal fitting is *α* = 0.45 (Fig. 3E). We used the point-to-plane distances between points and the projection plane to describe odor concentration variance of samples. Again, we plotted the radius-angle relationship with the sizes of points proportional to the variances (Fig. 3F). The inset shows that the variance increases with the spiral length of points. The results from Figure 3 show that the ripeness of strawberry progresses in a spiral shape and the variances of odor concentration profiles increase with the ripeness stage.

**Figure 3:**
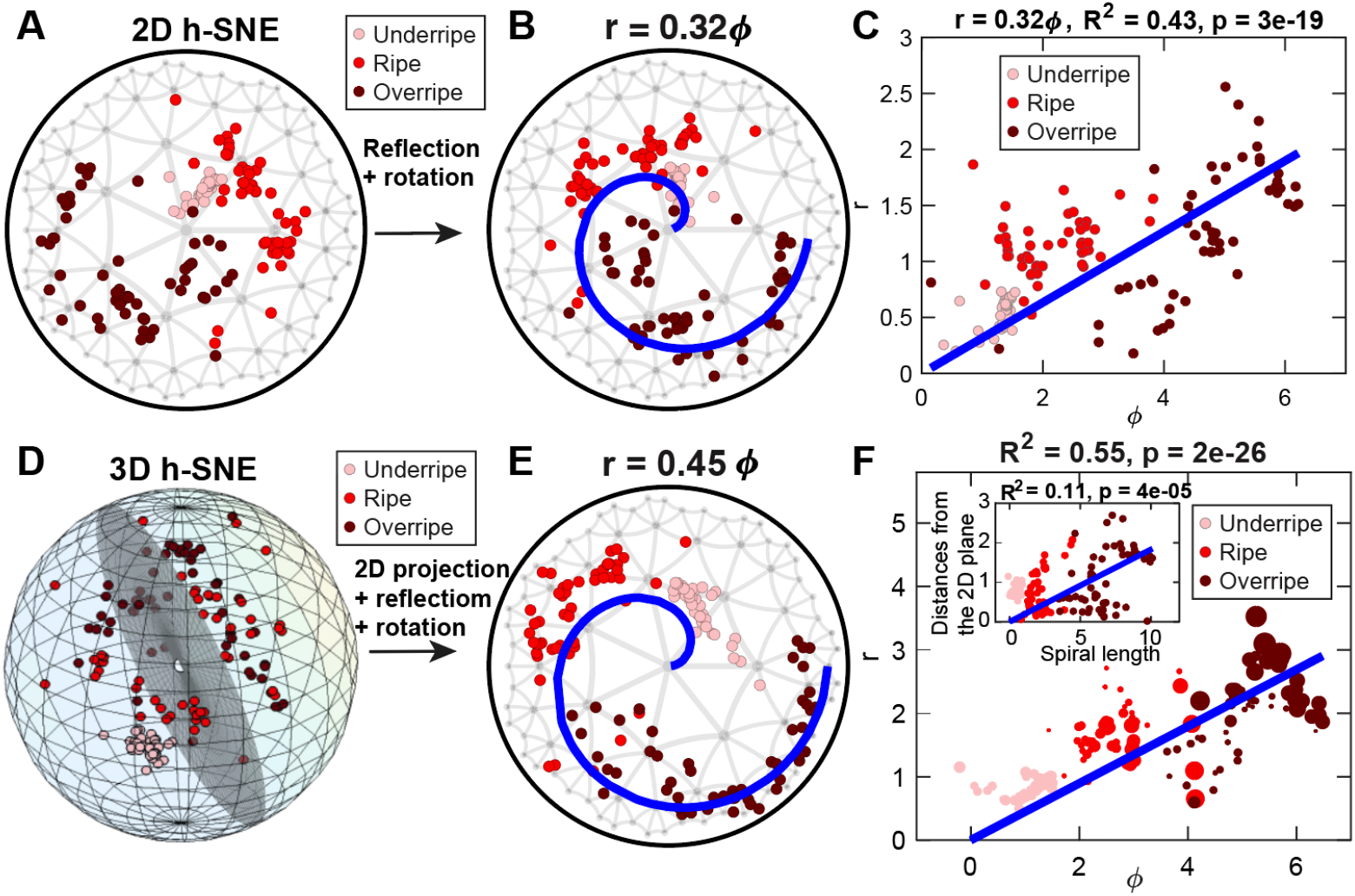
h-SNE embedding and spiral fitting of strawberry data. **(A)** h-SNE embedding of strawberry samples. The pink, red and dark red points represent under-ripe, ripe and overripe strawberries. **(B)** Invariant transformation of points in (A) by reflection and rotation, such that the points can best be fitted by an Archimedean spiral (blue line): *r* = *αϕ*. **(C)** Plot of radial coordinate of points in (B) as the function of the angular coordinate; the blue line is the linear fitting. **(D)** 3D h-SNE embedding of strawberry with a 2D intersection plane shown in gray, where the projected points have optimal spiral fitting after reflection and rotation. **(E)** Spiral fitting of projected points in the 2D disk shown in (D). **(F)** Plot of radial coordinate of points in (E) as a function of the angular coordinate. Sizes of points represent the point-to-plane distances of points in (D), and the inset shows the point-to-plane distances versus corresponding spiral lengths of points in (E).

### Collective spiral progression of multiple fruits in the fermentation dataset

Next, we studied the progression of the fermentation trajectory of the five types of fruits in the fermentation dataset (banana, kiwi, mango, peach, strawberry). Fruits undergoing uncontrolled or yeast fermentation were sampled on successive fermentation days, and the data were collected from between one to three different runs. The hyperbolic t-SNE embedding exhibited batch effects across runs that amounted to rotation of the data (Fig. 4A, left). To compensate for these effects and combine data across runs, we performed Procrustes analysis to align data across runs (Fig. 4A, right, see Methods). After batch correction, the five fruits collectively formed a spiral shape in the disk (Fig. 4B, left). We then performed reflection, orthogonal rotation, and translation on the embedding points, while preserving the pairwise distances, so that the points were best fitted by a two-parameter Archimedean spiral:

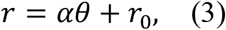

where *α* describes how fast the spiral expands with the angle and *r*_0_is a constant representing the initial radius of the fermentation. The optimal fitted parameters are: *α* = 0.59, *r*_0_ = 0.50 (Fig.4B, right). The linear relationship between radii and angles is shown in Fig. 4C. We further studied how the fermentation time was related to the spiral lengths of the points. We removed outlier samples from fermentation days of 10 and 17 for two reasons: first, these two days have a smaller number of samples (6 samples for each) compared with other days and they were only available in Run 1. After removing the outliers, the mean spiral lengths of the samples were well fitted by logistic function:

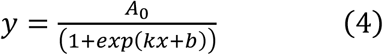

where *A*_0_ is a constant representing the upper bound of the spiral length. We set *A*_*m*_ = 2.1 and got the two fitting parameters: *k* = −0.11, *b* = −0.66 (Fig. 4D).

**Figure 4:**
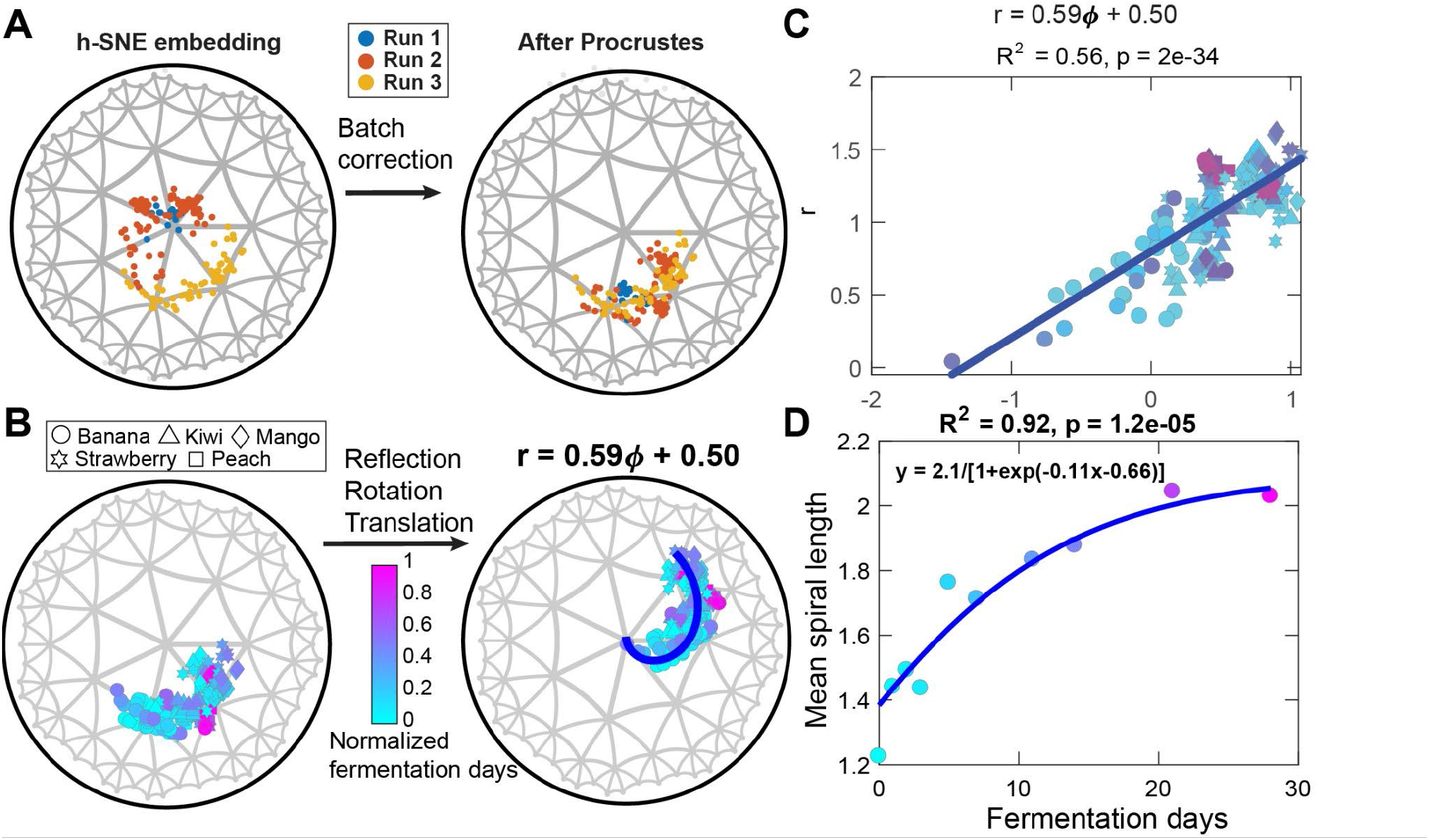
h-SNE embedding and spiral fitting of fermentation data. **(A)** h-SNE embedding of integrated fruit fermentation datasets from three runs (left) and batch correction by Procrustes for the three runs (right). **(B)** The batch-corrected samples are color-coded according to fermentation day and symbol-coded according to fruit type (left). The optimal Archimedean spiral fitting after performing distance-invariant reflection, rotation, and translation operations (right). **(C)** Plot of radial coordinate of points in (B) as a function of the angular coordinate. The blue line shows the linear fitting, and the color scales are the same as in (B). **(D)** Plot of the mean spiral length of samples as a function of fermentation day. Samples at fermentation day 10 and 17 days were removed as outliers (see text). The points are fit by a logistic function with a manually assigned amplitude (Eq. 4).

### Controlled fermentation yields more constrained spiral progression

To study how yeast affects the fermentation progression, we separated the uncontrolled fermentation and yeast fermentation conditions and fit them separately (Fig. 5A-B). We observed no significant difference in parameter *α* (0.65 for uncontrolled and 0.69 for yeast); however, the parameter *b* differed significantly with *a* = 0.54 ± 0.08 for uncontrolled and *a* = 0.31 ± 0.13 for yeast (Fig. 5A-B). The difference is visualized in the integrated embedding in Fig. 5C, where points corresponding to controlled fermentation are more constrained with smaller radii. The contribution of yeast fermentation can be described by a vector (red arrow in Fig. 5C) that describes additional rotational shift under yeast fermentation (see Methods).

**Figure 5:**
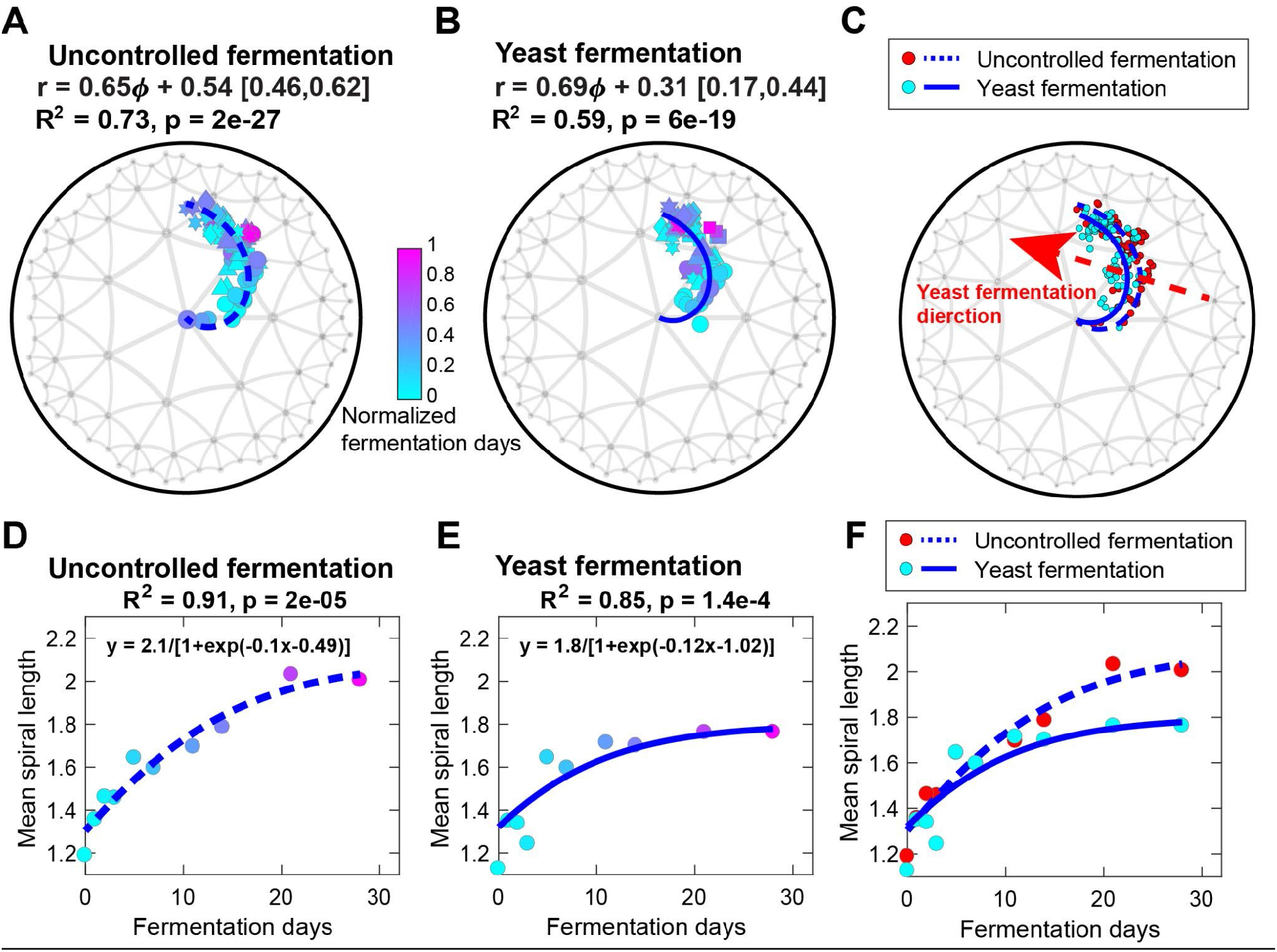
Comparison of yeast and uncontrolled fermentation odor mixtures. **(A-B)** Headspace profiles of uncontrolled (A) and yeast (B) fermentation were separately fit by Eq. 4. The 95% confidence interval of *b* is shown in the square bracket after the fitting equation. **(C)** Overlay of the two fittings in (A-B). The red arrow indicates the direction that yeast fermentation deviates from uncontrolled fermentation. **(D-E)** The same logistic fitting as in Fig. 4D for uncontrolled (D) and yeast (E) fermentation. Points are colored by fermentation day. **(F)** Overlay of the two fittings in (D-E).

Another way to analyze this shift is by “unfolding” the hyperbolic map and plotting radius vs. angle. In these coordinates, spirals correspond to straight lines and one can characterize the impact of various environmental factors, such as yeast, using linear methods. In this case, the uncontrolled fermentation map resulted in systematically larger radii compared to yeast fermentation (Fig. 5-supplemental), which could be interpreted as more rapid breakdown of complex molecules in the uncontrolled fermentation condition. This process could also be quantified by fitting the mean spiral length versus fermentation days with a logistic function as in Fig. 4D. We observed that the uncontrolled fermentation has larger amplitude (*A*_0_ = 2.1) than yeast fermentation (*A*_*0*_ = 1.8). These results show that controlled fermentation is more constrained in the progression in hyperbolic space, corresponding to faster angular rotation of the spiral.

### Identification of odors that drive ripeness and the fermentation process of fruits

We next asked how individual odors contribute to the orderly progression of fruit odor mixtures along the spiral in the embedding space. To make cross-dataset comparisons, we focused on the six common odors that are shared by the strawberry whole fruit dataset across stages and the multi-fruit fermentation homogenate dataset. We embedded the odor positions by minimizing the squared differences between sample positions and linear combinations of odor positions, weighted by the normalized odor concentrations (see Methods). The embedded odors are shown together with the samples in the same space and colored by the Pearson correlation coefficient of odor concentration with spiral length (Fig. 6A). The odor with the highest correlation of concentration to spiral length (styrene), i.e. with the largest contribution to ripeness, is located far from the odor with the smallest correlation (1-hexanol) (Fig. 6A). This relationship was also true for the fermentation data (Fig. 6B). Thus, the relative positions of the embedded odors may represent the relative contributions of the odors to the ripeness or fermentation stages. In addition to the qualitative evaluation, the embedding of odors preserve the odor distances directly calculated from data, with *R*^2^ = 0.833 for strawberry data and *R*^2^ = 0.748 for fermentation data (Fig. 6C).

**Figure 6:**
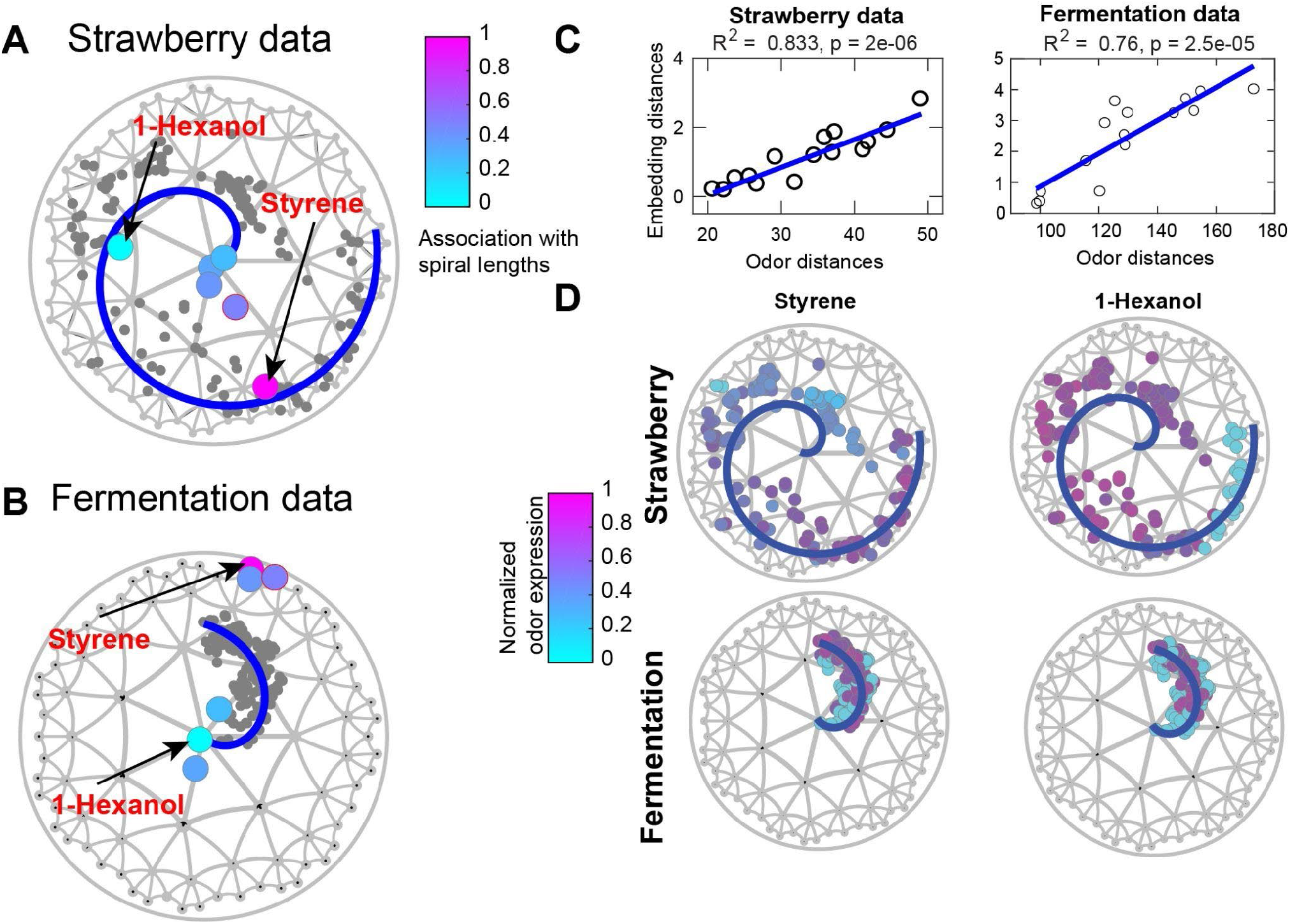
Identifying odors that drive spiral progression. **(A)** Visualizing odors together with natural mixtures from samples in the strawberry dataset in the same space (see Methods). The gray dots represent strawberry samples, and the colored dots represent six odors which are shared in strawberry and fermentation data. The color scale is determined by the Pearson correlation coefficient between odor concentration and spiral length in the strawberry data. **(B)** Visualizing the same six odors as in (A), together with natural mixtures from samples in the fermentation data. Points representing odors are color-coded the same as the corresponding odor in (A). **(C)** Pairwise distances of plots of odors in (A-B). **(D)** Abundance profiles of two odors styrene (left) and 1-hexanol (right) in strawberry (top) and fermentation (bottom) datasets. The color scales are determined by the normalized odor concentrations in each dataset.

From the above analysis, we identified two odors of interest: styrene with the highest association with spiral length and 1-hexanol with the lowest association. These two odors indeed showed distinct concentration patterns across samples. The concentration of styrene consistently increases with spiral length in both strawberry and fermentation data (left column in Fig. 6D), consistent with this odor being an indicator of fruit over-ripeness. Another odor that consistently increases with fermentation is ethyl butyrate. However, the concentration of 1-hexanol peaks at an intermediate stage in both datasets (right column in Fig. 6D).

### Estimating odor source phenotypes from volatile profiles collected at different stages of ripeness or fermentation

We focused this study on fruits in varying stages of ripeness and fermentation because we predict that fruit in these different states will have different behavioral meaning to animals (e.g., approach and taste, or avoid). To estimate the behavioral value of the natural odor mixture profiles we measured, we made use of a previously published dataset from Schwieterman et al.^28^ which measured behaviorally important properties of different strawberry samples, such as sweetness or sugar content, along with the odor profiles of each sample. We first calculated the association of odorant compounds with different phenotypes in this dataset, and then used our own measurements of volatile concentrations in the headspace profiles of our fruit sources in different states to assign a weight for its contribution to each phenotype. The integrated value across odorants in the mixture was calculated as an estimate of the phenotype for a given sample (see Methods). There are 11 available phenotypes in the published Schwieterman et al dataset^28^. We first performed hierarchical bi-clustering to these phenotypes and selected three phenotypes that have distinct patterns across the samples: sourness, malic acid and sweetness (Fig. 7A). We then displayed the patterns of the three phenotypes by coloring the points in 2D h-SNE map, according to the estimated phenotype values for samples in both the strawberry (top row) and fermentation (bottom row) datasets (Fig. 7B); the three phenotypes show distinct patterns along the spiral in both datasets. In order to visualize the patterns in a more quantitative way, we plotted the phenotype value as a function of spiral length (Fig. 7C). Considering samples progressively along the spiral, their predicted sweetness decreases, predicted sourness increases slightly, and their predicted malic acid content increases during the fermentation process (Fig. 7C, bottom row). In the strawberry dataset, these phenotypes show more complex patterns from under-ripe to over-ripe stages (Fig. 7C, top row). However, the strawberry ripening dataset represents a broader range of metabolic states, and the fermentation process that dominates the changes in the dataset may correspond to an intermediate stage in the strawberry dataset as the samples first transition from ripe to rotting/fermenting. Therefore, we selected a shaded area in the strawberry ripeness process in which the phenotype patterns would be predicted to best correspond to the fermentation process (Fig. 7C, top row). There results show that olfactory signatures produce expected changes in behavioral rankings that correspond with increased sourness and malic acidity and decreased sweetness as the fermentation process proceeds beyond the initial ripe stage.

**Figure 7:**
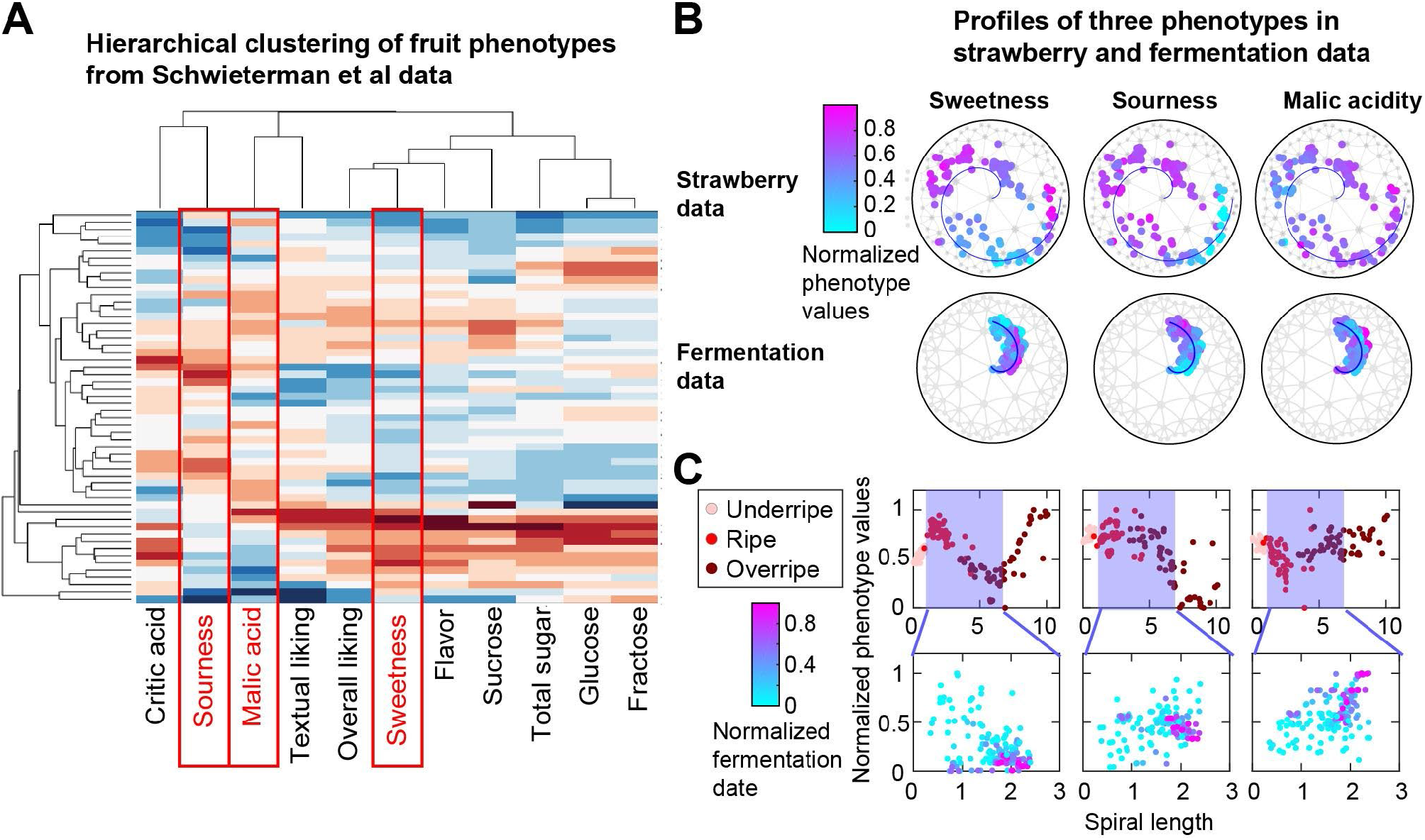
Estimation of odor source phenotypes for odor profiles in strawberry and fermentation datasets. **(A)** Hierarchical clustering of strawberry phenotypes based on concentration measurements of strawberry headspace volatiles from Schwieterman et al^28^. The rows and columns represent samples and phenotypes, respectively. The red boxes highlight three distinct phenotypes: sourness, malic acid, and sweetness. **(B)** Estimated sweetness, malic acidity, and sourness values of strawberry samples (top) and fermentation fruit samples (bottom) represented in h-SNE map. **(C)** Plots of estimated phenotype values versus the spiral length of points from strawberry samples (top) and fermentation fruits samples (bottom). The points are colored according to the ripeness stage for strawberry data and fermentation day for fermentation data. The blue shaded areas in the top panels correspond to the transition from ripe to rotting/fermenting in the strawberry dataset, which may best correspond to the fermentation process in the fermentation dataset (bottom). Sweetness decreases, whereas sourness and malic acidity both increase during fermentation that starts from ripe fruit.

## Discussion

This study demonstrates that plant and fruit volatiles can be organized according to a low-dimensional hyperbolic map, and that within this map exist trajectories that correspond to important metabolic processes, such as ripening or fermentation driven by the presence of microorganisms such as yeast.

The curved nature of odor space that we have described here and elsewhere^37,44^ arises naturally from the metabolic relationships in the pathways that produce odors^45-47^. The metabolic pathway have hierarchical organization ^48^, and as a result, the mixtures that they produce are expected to have a low-dimensional hyperbolic geometry. In a hyperbolic map, points close to the center describe “parent” molecules, while those further from the center describe derivative products of the initial reaction. A straightforward degradation corresponds in this map to simple radial progression from the center to the periphery of the space. Here we observe a spiral, suggesting that degradation is accompanied by systematic shifts in the activity of different metabolic pathways (denoted by different angular positions). From this perspective, it is interesting to note that the participation of yeast in the fermentation process results in a systematic shift characterized by a yeast-specific vector in the hyperbolic space.

Our analyses of fruit ripeness states test the idea that important organization axes of odor spaces are specified by metabolic pathways^44,49^. The fruit odors that we analyze are complex mixtures of many different chemical components that arise from metabolic pathways within the fruits themselves or through production by other important organisms associated with the fruits. Different ripeness states, as we show, are represented by an overlapping but evolving set of odorants. Some odorants are common to all three ripeness states that we categorized, whereas others are specific to one or two. Subsets of these components could be produced by defined biochemical pathways or by rate-limiting enzymatic reactions that are programmed genetically. Thus, the occurrence of components within these subsets will be correlated to a greater or lesser extent depending on where they lie along metabolic pathways.

Furthermore, the evolving sets of odorants we describe likely reflect not just metabolic pathways of fruits but also pathways of bacteria and yeast as they begin to take hold on ripening fruits. Styrene, for example, arises from fungal metabolic pathways ^42^. Information about metabolic pathways present, and what state they are in, is used by different species of fruit flies, for example, that utilize fruits in different states for food and egg laying^38^.

Our analyses build further support for the emerging framework where topography of odor spaces is organized around metabolic pathways^44,49^. The initial suggestion for this came from observations of topography in hyperbolic spaces in odors produced by mouse and several species of fruit^44^. In the case of strawberry and tomato odorants the topography of the odor space matched the topography of human rankings for these fruit samples^44^. Second, Qian et al. tested the metabolism-based topographical organization of odor spaces^49^, by training a neural network model using human perceptual data and derived a Principal Odor Map that they then used to evaluate odorants in defined metabolic networks. The distance between pairs of odorants in any metabolic network correlated closely with distances in the odor map based on perceptual data.

Thus, odorant pairs that co-occur in natural odor sources were closer together in the odor map. Moreover, the sequences of reactions in a metabolic pathway predicted the pathways through the odor map. Because an odor map based on human perceptual data could predict metabolic relationships across datasets from several different, diverse animal species, the map could represent a fundamental and generalizable feature of odor spaces. Indeed, recent work in the fruit fly olfactory system indicates that neural representations of odor become increasingly organized around the relationships of odors in natural odor sources, relative to odor relationships defined by chemical structure^50^. Therefore, the emerging view from these studies is that olfaction has evolved to extract information about metabolic pathways important for animals as they seek essential resources in their environment.

## Methods

### Profiling of headspace volatiles from individual strawberry fruit

Strawberries used in this study were supplied by Good Farms (San Diego, CA) through regular shipments over a few weeks. All shipped strawberries were supplied in the unripe/green state. We collected headspace volatiles from strawberries at three ripening stages (ripe, underripe, and overripe) as follows: one pound of ∼25 medium-size strawberries were first arranged on a piece of white A4 paper and photographed for future reference, e.g. to determine the degree of their redness. Strawberries were then homogenized using a blender (Black and Decker Appliances) and poured into an odorless plastic oven bag (Reynold Kitchens, 406mm × 444mm). Volatile collection was performed using an electric air sampler (PAS-500, Spectrex, CA, USA). A Tenax adsorbent tube (henceforth filter, Gerstel) was connected to the air sampler through a 20 cm Teflon tube. After the filter end inside the bag was brought into proximity with the blended strawberries, the bag was sealed. The headspace air was pulled at ∼ 100 ml/min through the filter which trapped the volatiles. After ∼4 hours of volatile collection, the filter was removed and transferred to a chemical laboratory for analysis of its contents according to the procedure described below. For the overripe stage, we set aside a portion of the ripe strawberries for ∼ one week until they became overripe, and then ran the volatile collection procedure described above. For each ripeness stage we collected 40-56 samples. Since we had the equipment to run 7 samples and 1 control each day we executed the sampling within ∼20 consecutive days.

For the chemical analyses, the filter contents were injected into a gas chromatography (GC, Agilent 6890 N, Santa Clara, CA, USA) capillary column (DB-MS1, J&W Scientific capillary 30 m × 0.25 mm × 0.25 μm) using thermal desorption technology (Gerstel) according to the following program: constant He flow (1 ml/min), column oven initial temperature was set at 40°C for 2 min, increased 7°C/min to 150°C followed by 20°C/min ramp to a final temperature of 250°C for 5 min. Identification and quantification of the strawberry headspace volatiles was performed using the GC connected to an Agilent 5975 mass spectrometer (GC-MS).

### Profiling of headspace volatiles from fruit in uncontrolled and yeast fermentation states

Ripe strawberry, mango, kiwi, banana, and peach in a state desirable for human consumption were acquired from local grocery stores. For each fruit, we measured the headspace profile of samples treated in two ways, yeast fermentation and uncontrolled fermentation. Both types of samples comprised 30 mL of homogenized peeled (except strawberry), unwashed fruit, added to a sanitized fermentation vessel, which consisted of a 200 mL mason jar with a fermentation airlock (Sauer system). To the yeast fermentation samples, 500 mg of yeast nutrients and 50 mg of Campden tablet, dissolved in 1mL of water, were immediately added to kill existing bacteria and inhibit wild yeast growth. Twenty-four hours later, 100 mg of wine yeast (Red Star) suspended in 1 mL of room temperature water was added to each yeast fermentation sample. Uncontrolled fermentation samples were left untreated and spoilage was driven by endogenous microbes present on the fruit. By 7 days of fermentation, uncontrolled fermentation samples appeared slimy and spoiled, sometimes with obvious fungal growth.

The volatile composition of the headspace of these fermentation vessels were sampled on days 0, 1, 2, 3, 4, 5, 11, 14 relative to the day that the sample was prepared. A few samples were additionally sampled at 21and 28 days after preparation. Three independent uncontrolled and yeast fermentation runs of banana, kiwi, and mango fruit were performed, starting from different fruit samples acquired on different days. towards the beginning of the fermentation. For peach or strawberry, only one or two, respectively, runs of uncontrolled and yeast fermentation were performed. To measure the volatile compound profile, we used solid phase microextraction coupled to gas chromatography-mass spectrometry (SPME/GC-MS), with a 75 μm CAR/PDMS SPME fiber (Supelco) thermally desorped into a Shimadzu GCMS-QP2020 single quadrupole GC-MS. The lids of the fermentation vessels had holes drilled into them which we covered with replaceable sanitized foil seals, through which we exposed the SPME fiber for 10 minutes. Fibers were desorped at 250°C for 1 minute, and samples were measured at splits of either 2 or 5, as needed to minimize detector overload. Oven temperature was programmed from 30°C (1 min hold) at 10°C/min to 150°C, then 60°C/min to 230°C (2 min hold). The GC used a ZB-5MS column (30 m length, 0.50 μm film thickness, 0.25mm i.d.), with a He carrier flowing at a linear rate of 52.1 cm/s. Compounds were identified using the Wiley (W9N11) and NIST/EPA/NIH (NIST 14) mass spectra libraries, with only matches of >95% similarity considered.

### Spiral length calculation

We calculate the spiral length of a point (*r*_*i*_, *θ*_*i*_) by the integration function:

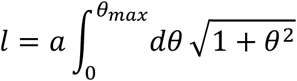

where *θ*_*max*_ = *θ*_*i*_ in strawberry data and 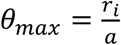 in fermentation data, *a* is the spiral coefficient in Eq. (1) or (3). There are two ways to calculate the spiral length from the position of a point: angle-based and radius-based. In the angle-based method, a given point is mapped to the point in the fitted spiral with the same angular coordinate, and this method is more reliable when the spiral is long and angular variation is small. In the radius-based method, a given point is mapped to the point with the same radial coordinate; this method is more reliable when the spiral is short and the radial variation is small. Based on these considerations, we apply the angle-based method on strawberry data because of the long spiral (Fig. 3B) and apply the radius-based method on fermentation data because of the short spiral (Fig. 4B). There is another bonus advantage for applying radius-based method to fermentation data: the integration parameter 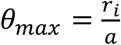 is independent of the second parameter *r*_0_ in Eq. (3) and invariant against rotation and reflection transformation. This property makes the spiral length calculation much easier.

### Batch correction using Procrustes algorithm

To correct the batch effects from three runs in fermentation data, we apply Procrustes to the embedding points. Procrustes determines a linear transformation (translation, reflection, orthogonal rotation, and scaling) of the points in a matrix Y to best conform them to the points in another matrix X, by minimizing the squared errors *L* between transformed matrices Z and X:

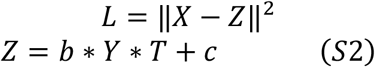

Where b is the scale component, T is the orthogonal rotation and reflection component, c is the translation component. Hence we can use Procrustes to align the samples in one batch to another reference batch. In the analysis in Fig. 4A, we use the samples in Run 3 as the reference batch and transform the samples in Run 1 and Run 2 by Procrustes. The three batches overlap with each other after the transformation (Fig. 4A).

### Embedding of odors together with samples

Given the positions of samples *P*_*sample*_, we infer the positions of odors *P*_*odor*_ by minimizing the squared errors between sample positions and combination of odor positions weighted by the normalized concentrations *C*. We take the following procedures:

1. Normalize each odor concentration *C*_*i*_ to be a unit vector.
2. *Minimize* (*P*_*sample*_ − *C* * *P*_*odor*_) ^2^, the optimal solution is: 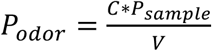, where *V* = *C* * *C*^*T*^.
3. Perform eigenvalue decomposition to *V*: *V* = *W* * *D* * *W*^*T*^,set eigenvalues below a threshold to be zero to reduce noise, then reconstruct a new matrix *V*_*thres*_ = *W* * *D*_*thres*_ * *W*^*T*^, and use *V*_*thres*_ to calculate *P*_*odor*_.
4. Uniformly rescale the radii of *P*_*odor*_ of all odors and calculate pairwise embedding distances.
5. Find the optimal threshold in step 3 and scaling factor in step 4 to maximize the Pearson correlation between pairwise distances of embedding points and data points.

As a result, 10 out of 13 eigenvalues are set to zero in strawberry data and 58 out of 61 eigenvalues are set to zero in fermentation data.

### Identifying yeast fermentation direction in tangent space

In order to find the yeast fermentation direction that contributes to the differences between sample positions from yeast and uncontrolled fermentation, we first transform the odor positions to tangent space by logarithmic map^51^:

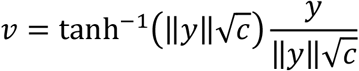

where c is the curvature of the space and is set to be 1 in our model. We calculate the mean differences of positions of samples from yeast and uncontrolled fermentation, then transform the calculated direction vector back to the original space by an exponential map:

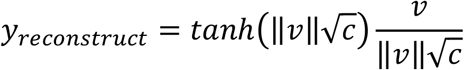

The final direction vector is shown as the red arrow in Fig. 5C.

### Estimating phenotype values

We estimate the phenotype values by taking the following steps (using sweetness as an example):

1. Calculate association of odor concentrations (*cct*) with sweetness intensity from Schwieterman data, use the Pearson correlation coefficient to define the sweetness of the odors: *sweet*_*odor* *i*_ = *ccrr*(*cct*_*i*_, *sweet*_*fruits*_)
2. Find the common odors between the new dataset and Schwieterman data.
3. Estimate fruit sweetness in the new dataset by the product of the common odor concentration and odor sweetness: *sweet*_*fruit* *k*_ = *cct*^*T*^ · *sweet*_*odors*_

## Data analysis tools

The violin plots were performed using R version 3.6.2, the other analyses were performed using MATLAB R2017a.

## Supplementary Figures

**Supplemental Table 1**. List of compounds in Fig. 1 Supplement 1. An “X” in columns E, F and G indicate components identified either in Fig. 1-Suppl Fig 2 or in the indicated references on analyses of strawberry odor components. Green, red and brown refer to sequential ripeness states in Fig. 1-Suppl Fig 1, whereas yellow indicates components also identified from yeast extracts. Finally, we include a list of odor and flavor descriptors for humans of all compounds identified in our chemical analyses.

**Supplemental Table 2**. List of compounds in Fig. 1 Supplement 2.

**Figure 1-Supplemental 1.**
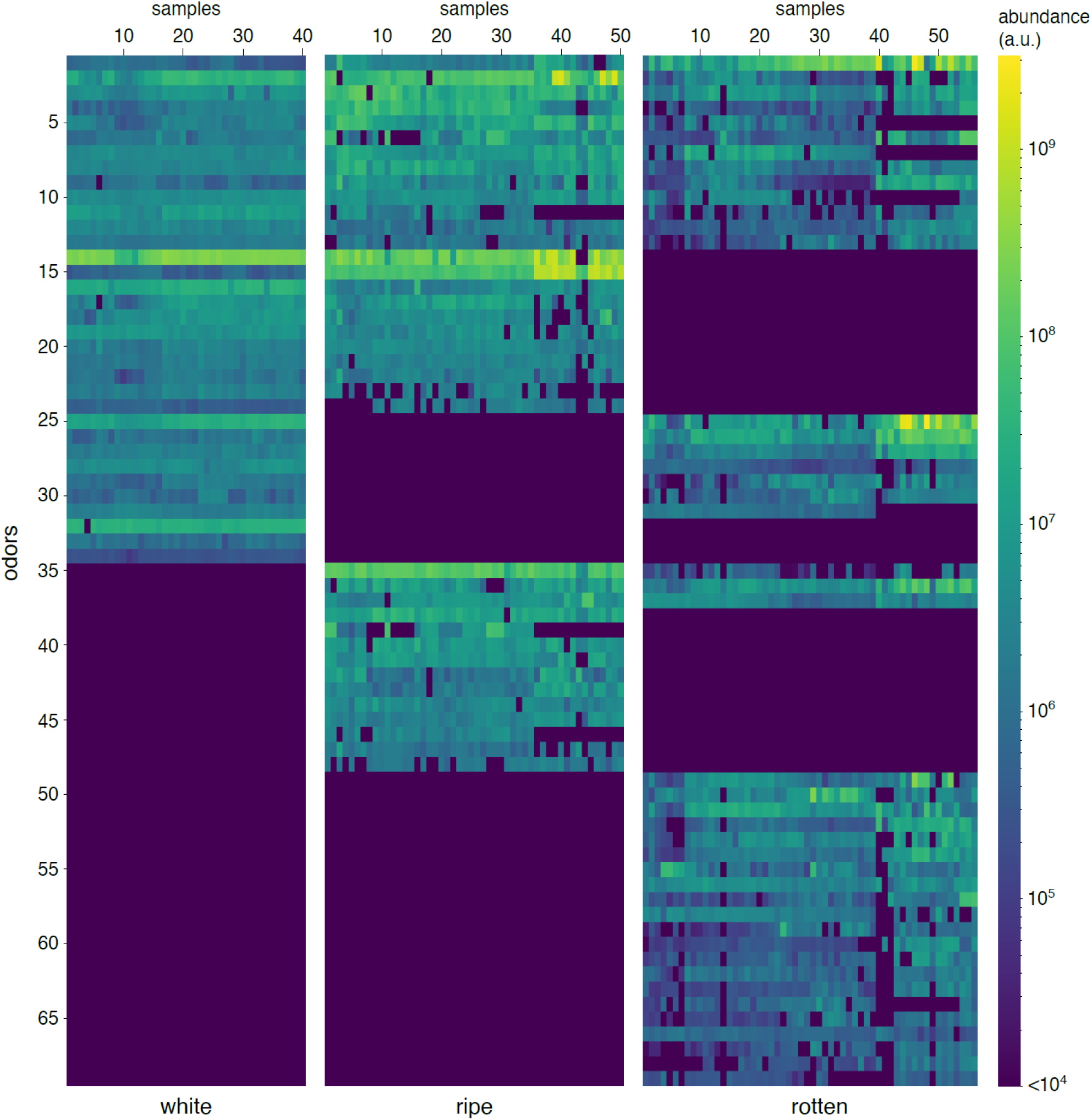
Compounds identified by dynamic headspace sampling from individual strawberry fruit in white (underripe), ripe, or brown/fermented states. Rows correspond to chemicals listed in the rows of Supplemental Table 1. Columns correspond to different individual samples. Color scale corresponds to relative abundance inferred from quantification of GC peak area.

**Figure 2-Supplemental 2.**
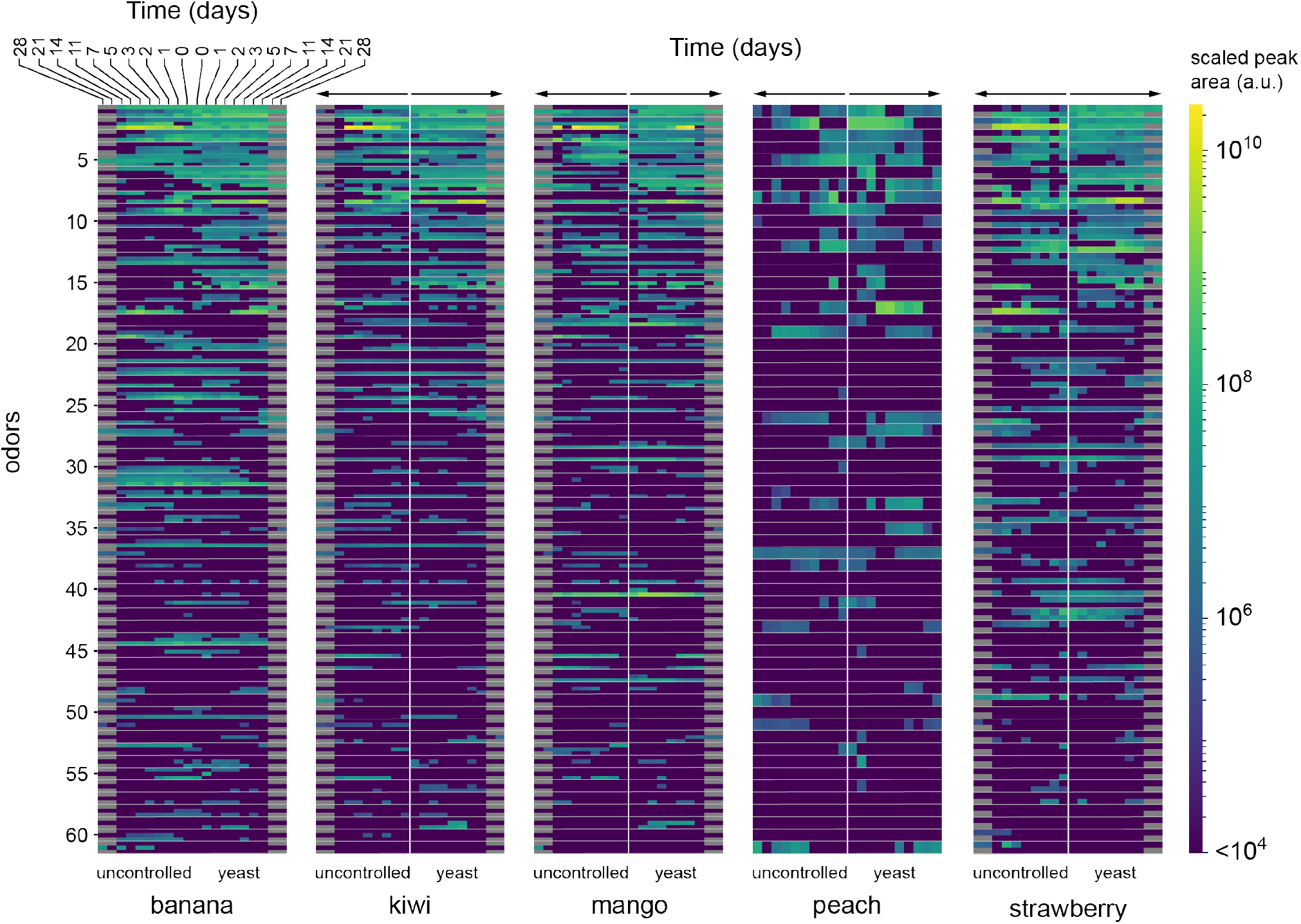
Compounds identified by static sampling (SPME/GC-MS) of the headspace of banana, kiwi, mango, peach and strawberry fruit mash undergoing uncontrolled or yeast-dominated fermentation. Fruits were sampled at 0, 1, 2, 3, 5, 7, 11, and 14 days of fermentation; some fermentation runs were additionally sampled at 21 and 28 days. Each row block corresponds to a different compound, specified in the rows of Supplemental Table 2. Each row block comprises 1-3 rows, corresponding to measurements from independent instances of the fermentation runs. Three independent banana, kiwi, and mango fermentation runs were measured, whereas only two independent runs of strawberry and one run of peach were measured. Columns correspond to different time points (in days) at which the samples were measured; the ordering of columns is the same for all five fruits. Color scale indicates the area under the curve for the peak corresponding to each compound in the gas chromatograph and indicates relative abundance.

**Figure 2-Supplemental.**
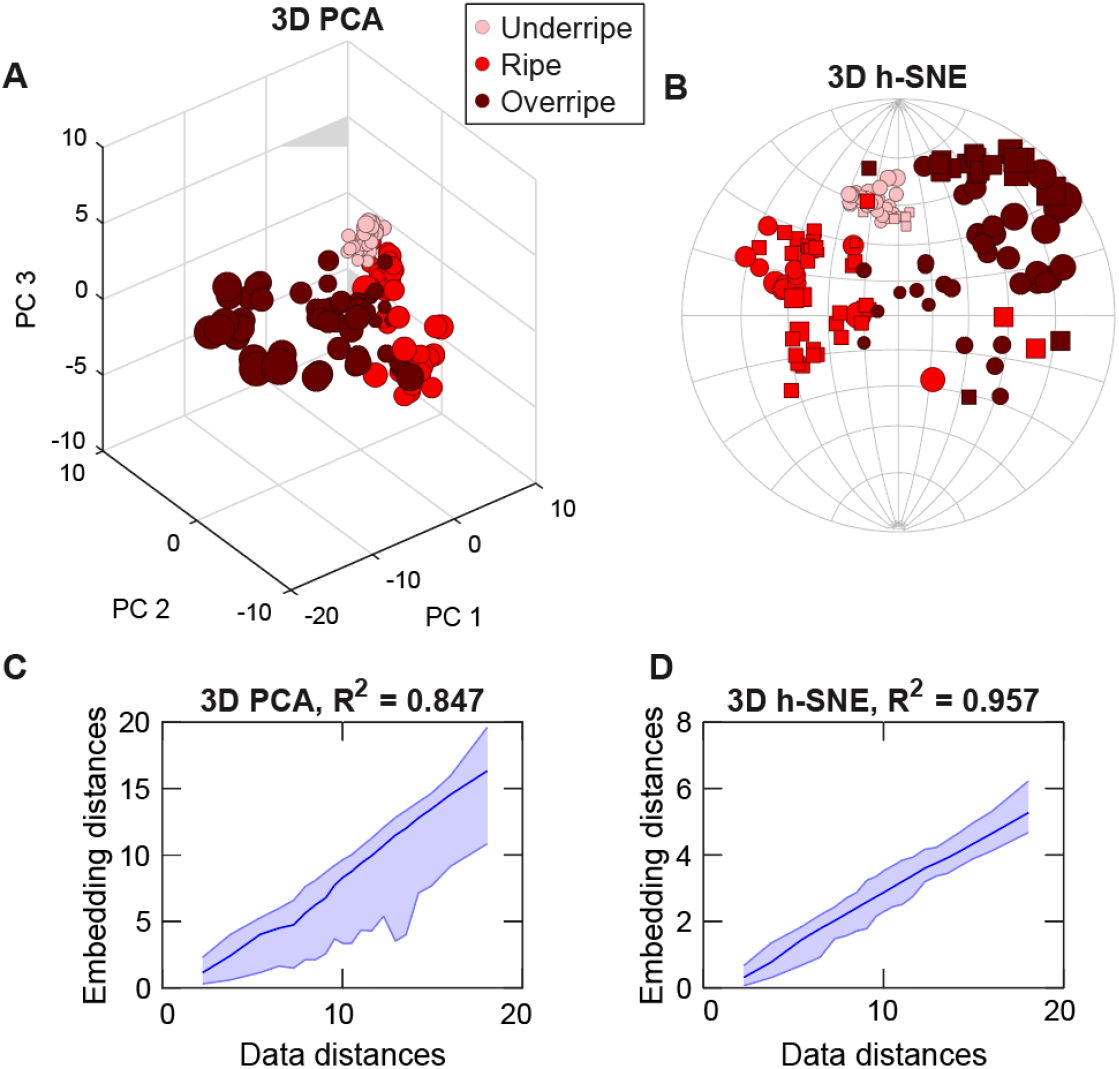
Embedding of strawberry samples by 3D PCA and 3D h-SNE. (A-B) 3D PCA(A) and h-SNE (B) embedding of strawberry samples with 13 common odors. (C-D) Distances preservation of PCA(C) and h-SNE (D) embedding.

**Figure 3-suppelemental.**
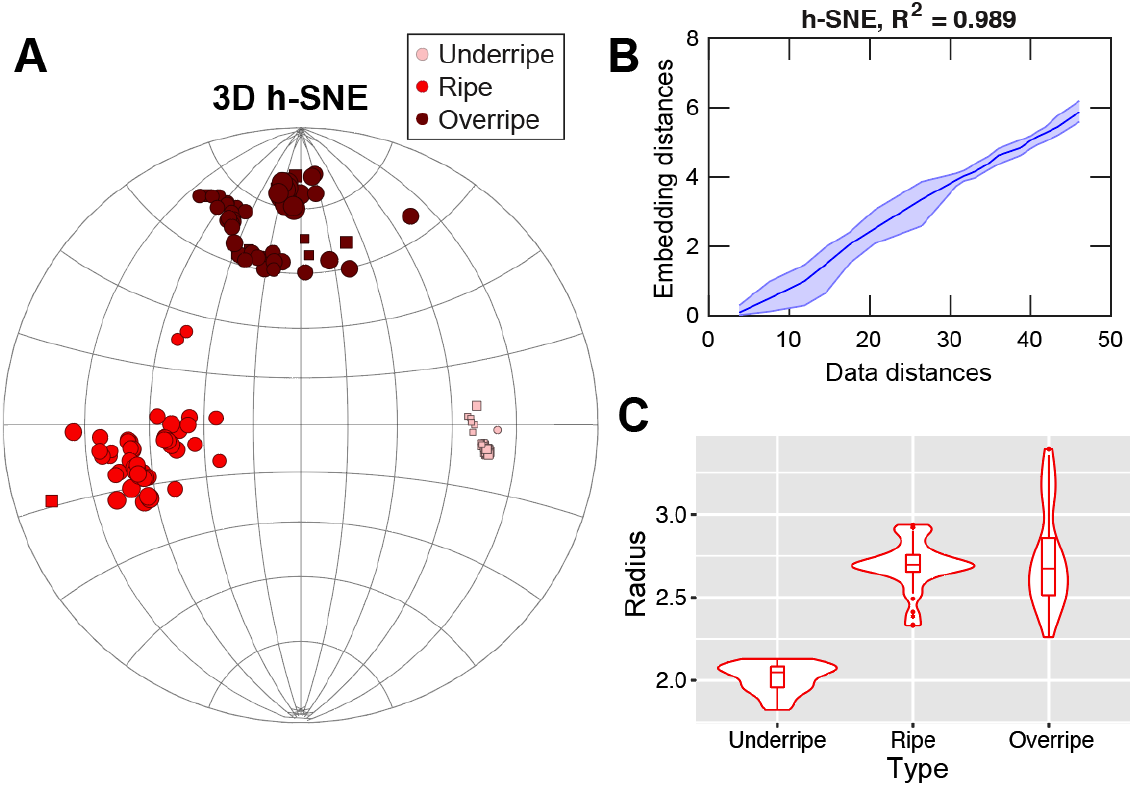
h-SNE embedding of strawberry samples by taking the union of all available odors. The missing values are padded by 100000. (A) 3D h-SNE embedding shown in a 2D stereographic plane. (B) Plot of embedding distances versus data distances. (C) Radii distribution of points in the three stages.

**Figure 5-supplemental.**
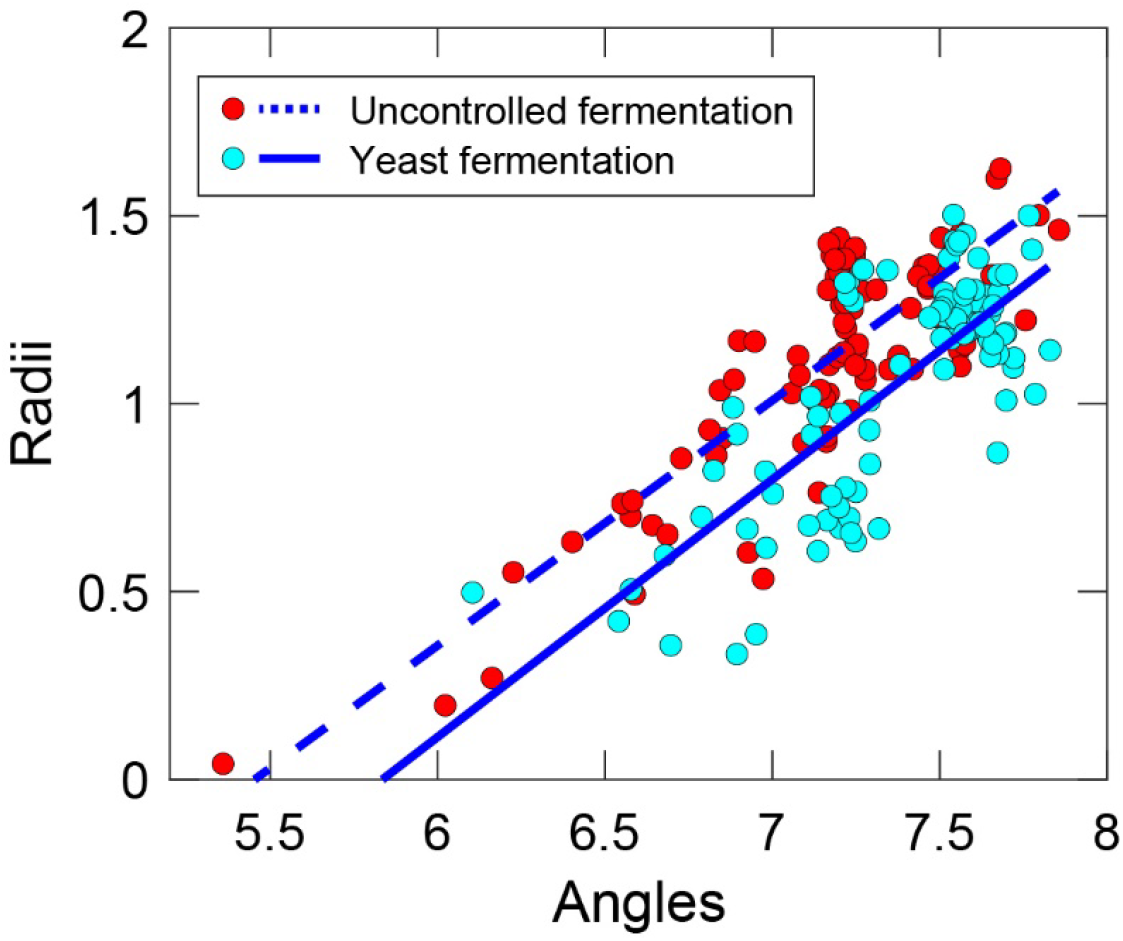
Effect of yeast fermentation in the radius vs. angle plane. Uncontrolled fermentation results in faster degradation represented by a larger radius of the fermentation spiral for the uncontrolled condition.

**Figure 6-supplemental.**
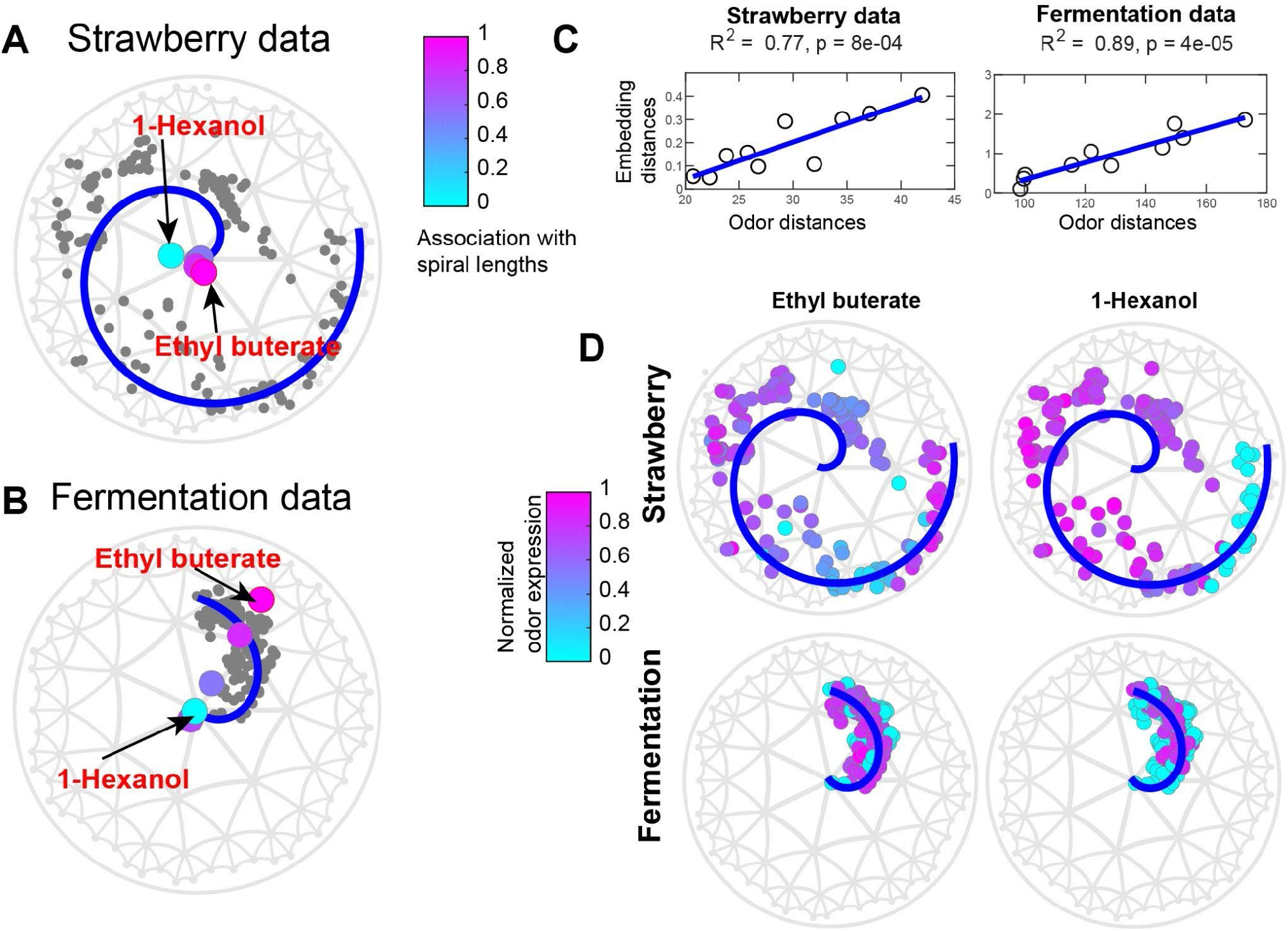
Analogous to Figure 6, but with styrene removed from both datasets. The results are similar to the case where styrene is included. In particular, the samples still follow a spiral trajectory (A,B) and correlation between fermentation date and spiral position is significant (B). (D) distribution of two example odorants in fermentation segments. Instead of styrene, we now highlight ethyl butyrate.

